# Characterization of clostridium botulinum neurotoxin serotype A (BoNT/A) and fibroblast growth factor receptor interactions using a novel receptor dimerization assay

**DOI:** 10.1101/718189

**Authors:** Nicholas G. James, Shiazah Malik, Bethany J. Sanstrum, Catherine Rheaume, Ron S. Broide, David M. Jameson, Amy Brideau-Andersen, Birgitte Jacky

## Abstract

Clostridium botulinum neurotoxin serotype A (BoNT/A) is a potent neurotoxin that also serves as an effective therapeutic for a variety of neuromuscular and glandular diseases and disorders. The observed pharmacological effect of BoNT/A is due to specific targeting and entry into motor nerve terminals within muscles followed by cleavage of the SNARE protein, SNAP-25, inducing a block in neurotransmission and temporary muscular paralysis. Specific binding and internalization of BoNT/A into neuronal cells are mediated by its binding domain (H_C_/A), which binds to GT1b ganglioside and protein cell surface receptors. Previously, fibroblast growth factor receptor 3 (FGFR3) was identified as a BoNT/A receptor, in addition to synaptic vesicle protein (SV2). To further study BoNT/A interactions with FGFRs, an FCS & TIRF receptor dimerization assay was developed to measure dimerization of FGFRs in live cells, as FGFR dimerization can be considered an indirect measure for receptor-ligand binding interaction and downstream signaling. The ability of H_C_/A to facilitate dimerization of three FGFR subtypes (FGFR1-3) was assessed. Recombinant H_C_/A (rH_C_/A) was shown to dimerize FGFR subtypes in the rank order FGFR3c > FGFR2b > FGFR1c. With potencies (EC_50_ values) defined as the concentration of ligand required to dimerize 50% of the receptors, wild type rH_C_/A dimerized FGFR3c with an EC_50_ of 24 nM, similar to FGF9, a native FGFR3c ligand, which had an EC_50_ of 15 nM, while FGFR1c and FGFR2b required higher rH_C_/A concentrations (≥100 nM and 68 nM, respectively). Furthermore, addition of the GT1b ganglioside to the culture media resulted in increased dimerization, whereas a ganglioside mutant variant of H_C_/A (rH_C_/A W1266L;Y1267S) showed decreased dimerization. Interestingly, reduced dimerization was also observed with an SV2 mutant variant of H_C/_A (rH_C_/A T1145A;T1146A). These results support a model wherein BoNT/A interacts with FGFRs, gangliosides, and SV2 on the cell surface to facilitate cell uptake.

## INTRODUCTION

Botulinum Neurotoxin Type A (BoNT/A) is a 150 kDa metalloenzyme belonging to the family of neurotoxins produced by *Clostridium botulinum* which causes temporary muscle paralysis by inhibiting acetylcholine release at the neuromuscular junction [1-4]. The neuronal specificity and high potency of BoNT/A has allowed its use in the treatment of a large number of medical and aesthetic conditions [3, 5-7], relying on injection of picomolar (pM) concentrations of the toxin. Though BoNT/A has been the subject of extensive study, greater understanding of the complex mechanism associated with BoNT/A’s neuronal specificity and cellular entry could lead to further therapeutic applications.

BoNT/A is a single-chain protein activated by proteolytic cleavage to form a 150 kDa di-chain molecule. The di-chain is comprised of a light chain (L_C_/A) which encodes a Zn^2+^ dependent endopeptidase (∼50 kDa), linked by a single disulfide bond and non-covalent interactions to a ∼100 kDa heavy chain (HC) containing the receptor binding and translocation domains [8]. The 50 kDa receptor binding domain, H_C_/A, is located at the C-terminal half of the HC and mediates specific binding and internalization of the toxin into neurons. Following internalization, the translocation domain (H_N_) of BoNT/A residing at the N-terminal half of the HC facilitates the translocation of L_C_/A from the endocytic vesicle into the cytosol. Once in the cytosol, L_C_/A enzymatically cleaves the soluble N-ethylmaleimide-sensitive factor attachment protein receptor (SNARE), synaptosomal-associated protein 25 (SNAP-25) [9, 10], which is essential for mediating vesicular fusion and exocytosis. Cleavage of SNAP-25 leads to inhibition of neurotransmitter/neuropeptide release, including acetylcholine, from neuronal cells and is responsible for BoNT/A’s observed pharmacological effects on smooth and skeletal muscles and glands [1, 11, 12].

Initially, BoNT/A binds with relatively low affinity (K_D_ ∼200 nM) to gangliosides, including GT1b, which are abundant at the presynaptic membrane and serve to trap and concentrate BoNT/A within the extracellular matrix [13-16]. H_C_/A mutations (W1266L;Y1267S)[16] have been identified which disrupt binding to GT1b and likely additional gangliosides (e.g., GD1a, GD1b, GQ1b, and GM1), which bind to BoNT/A with lower affinity compared to GT1b [14-17]. Once anchored close to the membrane, BoNT/A interacts with one or more relative high affinity protein receptors, including the synaptic vesicle protein, SV2 (K_D_ ∼100 nM) [18-23], and FGFR3 (K_D_ ∼15 nM) [24, 25]. Mutations within H_C_/A that disrupt binding to SV2 have been identified, including T1145A;T1146A [20], and G1292R [19]. The observed in vivo selectivity of BoNT/A for specific classes of neuronal cells is likely due to avidity upon binding to multiple receptors, which may also serve as a requirement to trigger internalization into the neuronal cell via endocytosis [13-16]. Interaction of BoNT/A with multiple receptors could provide an evolutionary advantage for the toxin, since it decreases the likelihood of host specific mutations resulting in toxin resistance. A similar strategy is known from other pathogens, including *Herpes simplex virus* (HSV) [26], *Trypanosoma cruzi* (Chaga’s disease) [27, 28], and human immunodeficiency virus (HIV) [29, 30]. Influenza A virus infection has also been suggested to involve FGFRs [31]. Previous protein complex immunoprecipitation results demonstrated an interaction between FGFRs and SV2 in neuronal cells [24], further suggesting the possibility of a multi-receptor BoNT/A complex. FGFR3 is one of four receptor-tyrosine kinases (FGFR1–4) that act as receptors for FGFs.

FGFR1–3, but not FGFR4, exist in three different splice variants, a–c, which differ in their extracellular ligand binding domains, each with differing ligand binding affinities and specificities [32, 33]. The a isoform variants terminate early, resulting in a secreted extracellular FGF-binding protein [34-37]. The b isoform variants, including FGFR2b, are primarily expressed in tissues of epithelial (surface tissue) origin, while the c isoform variants, including FGFR1c and FGFR3c, are primarily expressed in tissues of mesenchymal (connective tissue) origin. Native ligands for FGFRs are generally produced by either epithelial or mesenchymal cells and act on opposite tissue type FGFRs. An exception is FGF1, which binds to both b and c FGFR isoforms [38, 39]. There are 22 known FGF ligands that bind with different affinity and selectivity to the different FGFR splice variants. For example, FGF4 binds to FGFR1c > 2c > 3c, while FGF9 binds to FGFR3c > 2c > 1c and 3b, and FGF10 binds to FGFR2b > 1b [35, 37, 39, 40]. Most FGFs interact locally with FGFRs in a paracrine or autocrine manner, although a number of FGFs, including FGF19, FGF21, and FGF23, act like hormones in an endocrine manner [36, 41]. Selectivity and affinity in vivo is achieved via interactions with co-receptors, including: heparin, heparan sulfate (HS), neural cell adhesion molecule (NCAM), cadherin, integrin, Klotho, anosmin, neuropilin, and fibronectin, which interact with both FGFs and FGFRs [42, 43]. Gangliosides have also been reported to be co-receptors for FGFRs, affecting ligand binding, receptor dimerization, receptor activity, and subcellular localization [44, 45].

This tissue specific expression of the ligands, receptors, and co-receptors, guides tissue and organ growth and development. FGFR signaling upon ligand binding and receptor activation is associated with numerous cellular functions, including development, homeostasis, and metabolism. FGFRs are activated by dimerization induced by co-receptor and ligand binding, which enables the cytoplasmic kinase domains to transphosphorylate one another at specific tyrosine residues [34-37, 46].

As noted above, the binding domain of botulinum neurotoxin, rH_C_/A, has been shown to bind FGFR in vitro and in cells [47, 48]. To further study wild-type (wt) and mutant variants of rH_C_/A interactions with FGFRs and compare these to native FGF ligand interactions, a novel receptor dimerization cell-based assay was developed. Total internal reflection fluorescence (TIRF) microscopy was combined with fluorescence correlation spectroscopy (FCS) and number and brightness (N&B) analysis to develop an assay, hereafter referred to as the “FCS & TIRF receptor dimerization assay”, that measures receptor dimerization in live cells. By combining the two methods, the FCS & TIRF receptor dimerization assay allows monitoring of receptor dimerization at the cell membrane in real-time; in the presence of co-receptors, in any cell type, without the need to create a custom reporter cell line or reliance on the use of functional reporters, e.g. receptor phosphorylation sites or down-stream kinases.

In the study presented here, the FCS & TIRF receptor dimerization assay was used to evaluate FGFR dimerization in the presence of wt and mutant variants of rH_C_/A. FGFR dimerization with native FGFs possessing known receptor selectivity was evaluated in parallel. In addition, the impact of ganglioside and SV2 receptor interactions on H_C_/A mediated FGFR dimerization was studied. To better understand the specificity of the interaction between rH_C_/A and FGFRs, binding to another tyrosine kinase growth factor receptor, epidermal growth factor receptor (EGFR), was also evaluated. Our results demonstrate that rH_C_/A, in the presence of gangliosides and SV2, can induce dimerization of FGFRs with a preference towards FGFR3c, further supporting a functional role of FGFRs in BoNT/A cellular binding, as previously suggested [24].

## MATERIALS and METHODS

### Materials

All cell culture medium and transfection reagents were from ThermoFisher (Carlsbad, CA) unless otherwise stated. FGF proteins [2, 4, 13, 14] and EGF were from R&D Systems (Minneapolis, MN). Collagen IV coated glass bottom 35 mm imaging dishes were from MatTek (Ashland, MA). Rat pheochromocytoma cells (PC-12) and fetal bovine serum (FBS) were from ATCC (Manassas, VA). GT1b was from Enzo Life Sciences (Farmingdale, NY). C-HaloTag-FGFRs, which included FGFR1c (EX-Y2820-M50, accession number: NM_023110.2, Homo sapiens fibroblast growth factor receptor 1), FGFR2b (EX-Z6888-M50, accession number: NM_001144913.1, Homo sapiens fibroblast growth factor receptor 2), and FGFR3c (EX-M0098-M50, accession number: NM_000142.4, Homo sapiens fibroblast growth factor receptor 3), and C-HaloTag-EGFR (EX-A8661-M50, accession number: NM_005228.4, Homo sapiens epidermal growth factor receptor) constructs were from Genecopoeia (Rockville, MD). AlexaFluor488 Halo ligand (AF-488) was from Promega (Madison, WI). Protease inhibitor cocktail was from Sigma-Aldrich (St. Louis, MO). Benzonase nuclease and rLysozyme were from EMD Sigma (Billerica, MA). HisTRAP FF and DEAE columns were from GE Healthcare (Chicago, IL).

### Cell culture

PC-12 cells were cultured on 100 mm collagen IV plates (cat. no. 354453) using complete growth media consisting of RPMI media with 2 mM GlutaMAX™, 10% FBS, 10 mM HEPES, 1 mM Sodium Pyruvate, 100 U/ml Penicillin, and 100 µg/ml Streptomycin. Cells were seeded onto imaging dishes and incubated in complete medium supplemented with 25 μg/mL GT1b (unless otherwise noted) until reaching confluency (∼4 days). The cells were maintained at 37°C with 5% CO_2_.

### Transfection of cells

PC-12 cells were plated at ∼1×10^6^ cells per 35 mm collagen IV coated glass bottom imaging dish and transfected with a mammalian expression plasmid containing the DNA sequence encoding C-HaloTag-FGFR1c, C-HaloTag-FGFR2b, C-HaloTag-FGFR3c, or C-HaloTag-EGFR. A DNA solution was prepared by adding and mixing (by inversion) 2.5 μg of plasmid with 100 µL OPTI-MEM media. The transfection solution was prepared by adding and mixing (by inversion) 5.0 μL of Lipofectamine 2000 with 100 µL of OPTI-MEM media. Once the DNA and transfection solutions were mixed by inversion and incubated at room temperature for 30 minutes, the 200 µL aliquot of the lipofectamine/plasmid solution was added to the plate containing the cells. After 6 hours of incubation, the medium was replaced with complete medium with or without GT1b and incubated for an additional 24 hours prior to addition of the HaloTag ligand.

### Treatment of cells with Halotag ligand and test compound for cell imaging

After transfection, the cells were incubated with 500 nM AlexaFluor488 Halo ligand for 15 hours. The cells were then washed three times (10 minutes/wash) with fresh complete medium to remove free ligand and incubated in serum-free medium (complete medium minus FBS, plus B-27 and N-2) for 3-4 hours prior to imaging.

### Expression and purification of wildtype and mutant variants of H_C_/A

Recombinant H_C_/A (rH_C_/A; spanning aa 876-1296 of full length BoNT/A1; GenBank accession no. AF48874) was cloned into pET28a+ (N-terminal His6-tag). The DNA was codon optimized for expression in *Escherichia Coli*. The variants of H_C_/A, (W1266L;Y1267S [16]) and H_C_/A (T1145A;T1146A [20]) were made by site-directed mutagenesis (Genewiz, NJ). The variants were expressed and purified as previously described [24]. Briefly, chemically competent BL21(DE3) *E. coli* (Thermo Fisher Scientific, Waltham, MA) were transformed with a rH_C_/A expression plasmid. Cells were grown at 37°C until OD600 reached 0.7 and expression induced with 1 mM Isopropyl β-D-1-thiogalactopyranoside (IPTG). After 16 hours of growth at 22°C, cells were harvested via centrifugation and lysed in buffer containing: 50 mM Tris-HCl pH 8.0, 10 mM EDTA, 100 mM NaCl, 10 mM DTT, 5% (v/v) glycerol, cOmplete™ EDTA-Free Protease Inhibitor Cocktail (MilliporeSigma, St. Louis, MO), 150 mU/mL rLysozyme, and 50 mU/mL benzonase nuclease, and then sonicated for 5 minutes. Lysate was cleared through high speed centrifugation and subjected to Immobilized Metal Ion Affinity Chromatography (IMAC) (HisTrap™), buffer exchange (desalting column) and Ion Exchange Chromatography (IEX) using the ÄKTAxpress system (GE Healthcare Bio-Sciences, Pittsburgh, PA). Protein (50 mM HEPES, pH 7.4, 150 mM NaCl) was quantified using NanoDrop™ 2000 spectrophotometer (Thermo Fisher Scientific).

### Image acquisition

Images were recorded on a Nikon Eclipse Ti TIRF microscope using a 60X 1.45 NA TIRF oil objective. Approximately 1000 frames were collected per cell and captured on a cascade 512B EMCCD camera (Photometrics, Tucson, Az) at 100 frames per second using software written in-house [48, 49]. AF-488 was excited at 488 nm (Sapphire SF 488; Coherent, Santa Clara, CA) with less than 30% laser power using an excitation filter cube (488/10 nm bandpass and 500 nm long pass emission filter; Chroma, Bellows Falls, VT) within the infinity space. Intensity images were recorded for each transfected HaloTag receptor in the absence of FGF or rH_C_/A protein ligand at both the beginning and end of each experiment from two separate dishes. The cells were then treated for 30 minutes with increasing concentration of ligand, reconstituted in PBS containing 0.1% Bovine Serum Albumin at a final concentration of 0.1 to 200 nM, prior to recording TIRF intensity images. Dark counts for the EMCCD camera were recorded at the beginning and end of each experiment. Free AF-488 (90 nM) in serum-free medium was used as the monomeric standard. Images were processed using the N&B function on the software platform SimFCS. Using FCS, the brightness of a particle as well as the number of particles in a given volume (Number and Brightness Analysis, S1) were separately obtained to determine the degree of aggregation of proteins in solution [47-53].

The average normalized brightness value ± standard deviation (SD) for each concentration of ligand was calculated based on at least 20 individual cells on 4 independent days. The results were plotted using GraphPad 7.02 Prism and fitted to a two- or three-parameter logistic curve (2PL or 3PL) using the log of the concentration to estimate the EC50 values and the 95% confidence intervals (95% CI). For the fitting, the curves were constrained with bottom >1 (3PL) or =1 (2PL) and the top =2, reflecting a transition from monomer to dimer state. For graphical presentation, the concentrations are plotted on a log scale.

## RESULTS

### FCS & TIRF receptor dimerization assay for studying receptor dimerization in live cells

In the TIRF (Total Internal Reflection Fluorescence) method, the optics of the instrument are adjusted such that the exciting light will be reflected from the interface, i.e., the glass surface supporting the cell. However, some of the energy of the incident beam will penetrate through the interface, creating what is termed an evanescent field which extends a very short distance on the order of 100 nm into the cell. Hence, this evanescent beam will only be able to excite fluorophores which are located near the cell surface—i.e., the plasma membrane of the target cell (**Figure 1**) [54]. Fluorescence fluctuation spectroscopy and number and brightness (N&B) analysis is an emerging method for analysis of molecular interactions in living cells and is based on the statistical analysis of signal fluctuations emitted by fluorescently labeled molecules [47-57] (**S1 Figure**). The method utilizes the fact that fluorescently labeled proteins produce intensity fluctuations as they pass through a small observation volume. Since the method relies on movement of fluorescent molecules, only mobile molecules are evaluated. While the average intensity in two sample volumes containing the same number of fluorophores may be the same, the fluctuations in intensity depend on the molecular brightness of the fluorescent protein molecules in the samples, and the magnitude of the fluctuations contains information about molecule concentration and the oligomeric state. By measuring the average fluorescence intensity within each pixel (photon counts per second per molecule), the average oligomeric state within each pixel is determined by calculating the molecular brightness, which is directly related to the stoichiometry of fluorophores in a protein complex, and normalizing it to a molecular brightness monomer standard. For example, if a fluorescently labeled monomeric FGFR is found to homodimerize under addition of a ligand, a 2x increase in the average molecular brightness would be observed compared to the monomeric brightness standard, and the number of pixels with 2x normalized average molecular brightness would increase (**Figure 2B; S2 Figure**).

**Fig 1.**
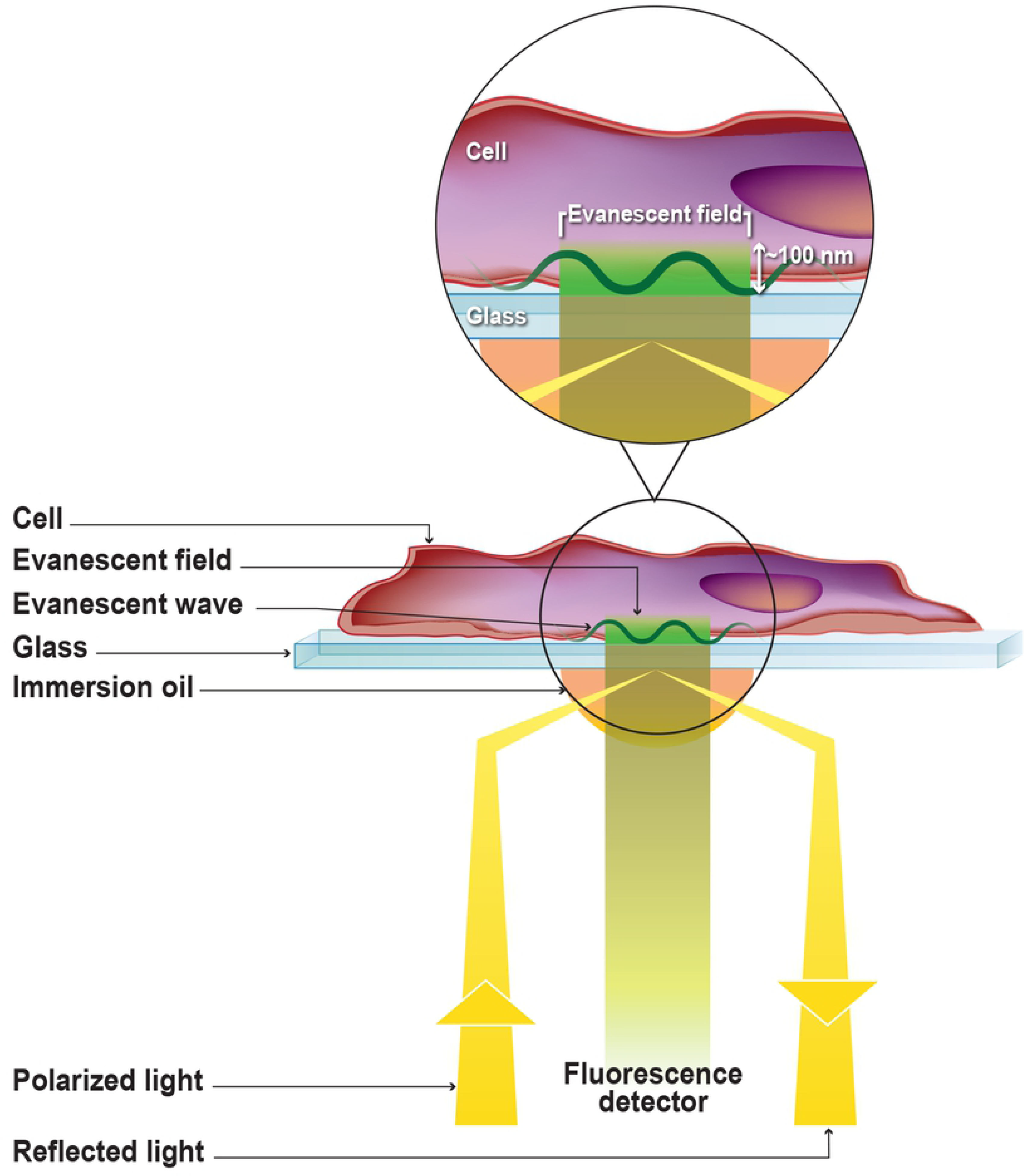
Illustration of the total internal reflection fluorescence (TIRF) method.

**Fig 2.**
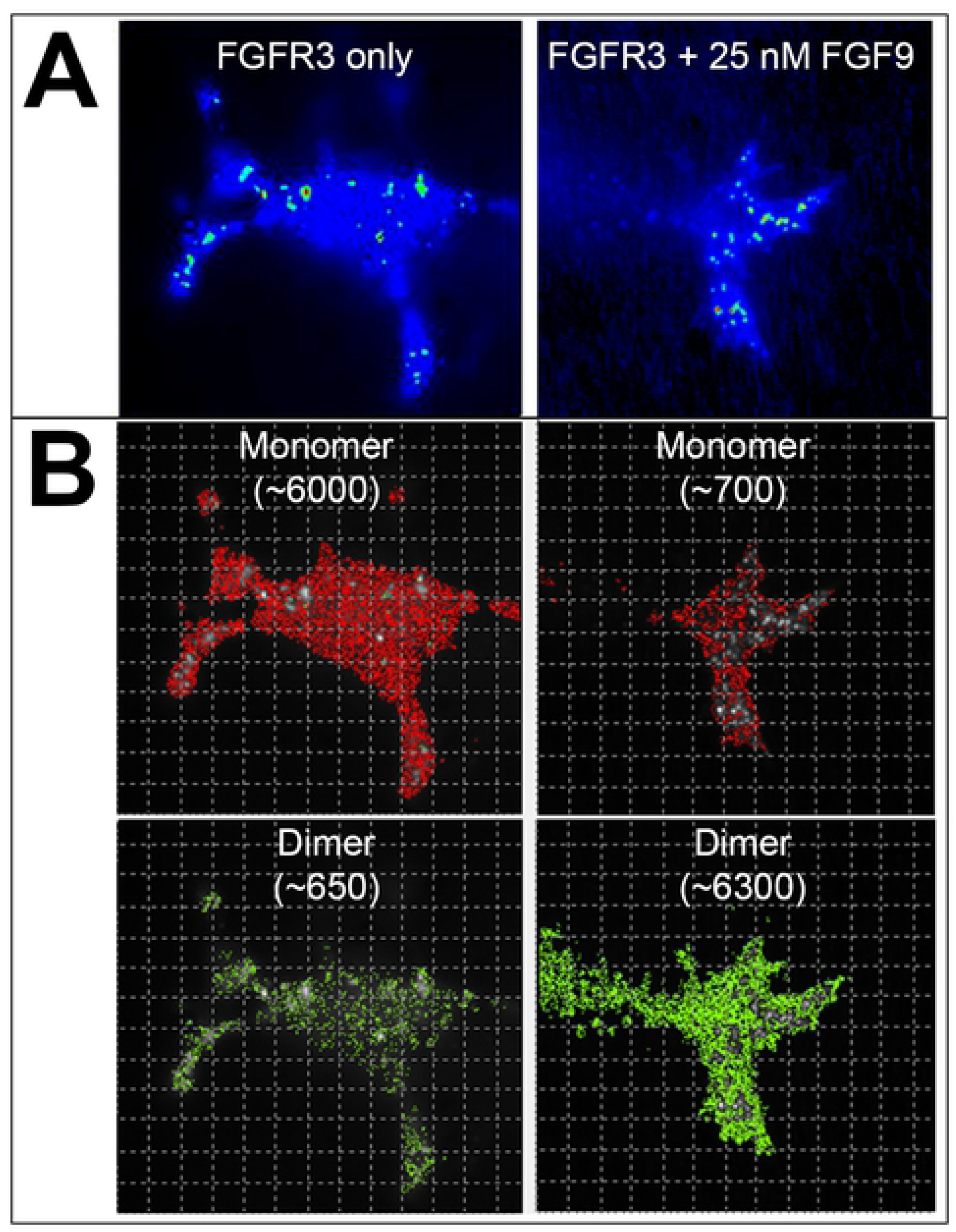
Depiction of intensity and molecular brightness images. **A.** (Intensity) Typical single cell intensity images are shown for cells that have not been exposed to FGF9 ligand (left column) and cells exposed to 25 nM FGF9 ligand (right column). **B.** (Molecular brightness) The normalized average molecular brightness images for each cell indicate monomer (red) and dimer (green) numbers and distribution throughout the cell with corresponding pixel counts, which is the number of pixels that contain dimer or monomer, respectively.

Using the FCS & TIRF receptor dimerization assay, the ratio of monomer to dimer FGFRs on the plasma membrane of neuronal PC-12 cells was measured, with and without addition of FGF ligand or rH_C_/A protein. We focused on FGFR1c, FGFR2b, and FGFR3c, which represents mesenchymal, epithelial, and neuronal FGFRs, respectively [32, 58, 59]. Briefly, a rat neuroendocrine tumor cell line, PC-12, was transiently transfected with FGFRs containing a C-terminal HaloTag, which is a modified haloalkane dehalogenase designed to covalently bind to exogenously added synthetic ligands bound to a fluorophore (HaloTag ligands; AF488 [cell impermeable for sell surface labeling]) [55] (FGFR3c-AF488). A dose dependent increase in ERK1/2 phosphorylation was observed in transfected versus non-transfected cells upon addition of native ligands, FGF2 and FGF9, at 2, 9, and 21 nM, indicating that the C-terminal HaloTag does not disrupt FGF receptor function (data not shown). TIRF intensity measurements of transfected PC-12 cells indicated a non-homogenous distribution of fluorescence, both with and without the ligand (**Figure 2A**), perhaps indicative of aggregated receptor proteins. Immobile fluorophores do not fluctuate, so any potentially aggregated protein will not be analyzed. Calculation of the average molecular brightness, in the absence of ligand, showed that the values for FGFR3c-AF488 were higher (1.1-1.2-fold) than the Alexa488 fluorophore molecular brightness monomer standard, suggesting that 10-20%of the receptors are in preformed dimers. Dimerization in the absence of ligand has previously been reported for EGFR under overexpression conditions that favor high receptor densities on the plasma membrane [60]. Data was collected from cells with an emission of 10,000-15,000 counts per cell, corresponding to approximately 100,000-150,000 receptors per cell, based on the Alexa488 fluorophore monomer standard. This criterion was used to ensure that the level of receptor expression was relatively constant across measured cells, avoiding cells with very low or high receptor densities. Typical intensity images corresponding to monomer and dimer populations are shown in **Figure 2**, where saturation with native ligand (FGF9 for FGFR3c) caused ∼2x increase in normalized average molecular brightness, consistent with transition to a 100% dimerized state.

### rH_C_/A induces FGFR3c dimerization

Previously, rH_C_/A was shown to bind to extracellular loops 2 and 3 of FGFR3 in vitro and on cells with a KD of ∼15 nM, which is similar to the affinity of native FGFR ligands, fibroblast growth factors (FGFs) [20]. To further validate the FCS & TIRF receptor dimerization assay and to compare the ability of FGFs and rH_C_/A to dimerize FGFR3c, PC-12 cells transfected with FGFR3c-AF488 were treated with rH_C_/A. In parallel, two native FGFR ligands were analyzed; FGF9, a known agonist ligand for FGFR3c and FGF10, a known agonist ligand for FGFR2b but not FGFR3c. As expected, addition of FGF9 resulted in a dose-dependent increase in average normalized molecular brightness, whereas treatment with FGF10 did not (**Figure 3**). EC_50_ values for FGFR3c dimerization, defined as concentration of ligand resulting in 50% receptor dimerization, were 18 nM (95% CI [12, 24]) for FGF9 and >200 nM for FGF10 (**Table 1**). Notably, the ability of rH_C_/A to dimerize FGFR3c was similar to FGF9 with an EC_50_ of 27 nM (95% CI; 18, 41), suggesting that rH_C_/A has FGFR3c agonist-like properties (**Figure 3; Table 1**).

**Table 1.** FCS&TIRF receptor dimerization assay – Result summary.

| Receptor | Ligand | GT1b | Function | EC<sub>50</sub> | 95% CI* |
| --- | --- | --- | --- | --- | --- |
| **FGFR1c** | FGF4 | Yes | Native FGFR1c Ligand | 17 nM | (8, 35) |
|  | FGF10 |  | Native FGFR2b Ligand | >100 nM | N/A |
|  | rH<sub>C</sub>/A |  | BoNT/A binding domain | 163 nM | (123, 203)** |
| **FGFR2b** | FGF10 |  | Native FGFR2b Ligand | 19 nM | (13, 26) |
|  | FGF4 |  | Native FGFR1c Ligand | >100 nM | N/A |
|  | rH<sub>C</sub>/A |  | BoNT/A binding domain | 70 nM | (59, 82) |
| **FGFR3c** | FGF9 |  | Native FGFR3c Ligand | 18 nM | (12, 24) |
|  | FGF10 |  | Native FGFR2b Ligand | >200 nM | N/A |
|  | rH<sub>C</sub>/A |  | BoNT/A binding domain | 27 nM | (18, 41) |
|  | rH<sub>C</sub>/A | No | BoNT/A binding domain | 44 nM | (36, 54) |
|  | rH<sub>C</sub>/A W1266L;Y1267S | Yes | BoNT/A binding domain, ganglioside mutant | 143 nM | (127, 161) |
|  | rH<sub>C</sub>/A T1145A;T1146A |  | BoNT/A binding domain, SV2 mutant | 78 nM | (53, 115) |
| **EGFR** | EGF |  | Native EGFR Ligand | 14 nM | (11, 19) |
|  | rH<sub>C</sub>/A |  | BoNT/A binding domain | >150 nM | N/A |

\*Based on 3PL with bottom >1, top =2
\*\*Based on 2PL with bottom = 1, top = 2

**Fig 3.**
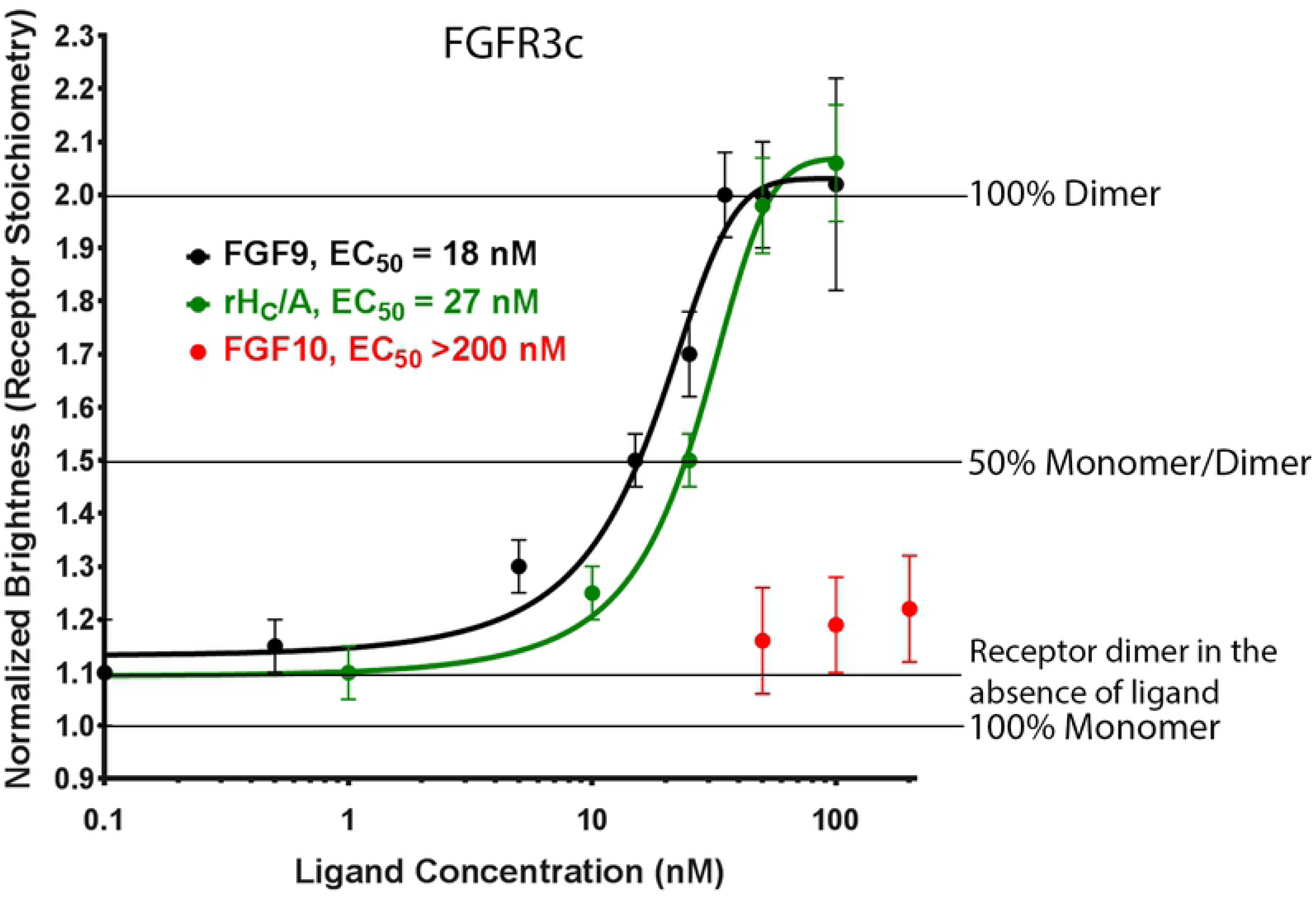
rH_C_/A induces FGFR3c dimerization. Native ligand FGF9 (black) (EC_50_ = 18 nM [95% CI; 12, 24]) and BoNT/A binding domain (rH_C_/A) (green) (EC_50_ = 27 nM [95% CI; 18, 41]), but not FGF10 (red) (EC_50_ >100 nM), dimerize FGFR3c expressed on neuronal PC-12 cells. Points represent the average normalized brightness values ±SD from greater than 30 cells collected on 4 independent days.

### rH_C_/A induced dimerization of FGFR3c is dependent upon GT1b ganglioside

Gangliosides, such as GT1b, serve as abundant, low affinity receptors for BoNT/A at the neuromuscular junction [13-16]. Since gangliosides can also function as co-receptors for FGFRs [44, 45], the FCS & TIRF receptor dimerization assay was performed with and without addition of exogenous GT1b to directly assess its potential role in rH_C_/A induced FGFR3c dimerization. A ganglioside mutant variant of H_C_/A (rH_C_/A W1266L;Y1267S [16]) was included to assess potential interactions with endogenous ganglioside produced in culture. Previously, it was shown that rH_C_/A W1266L;Y1267S protein lost its ability to bind to neuronal cells but maintained its ability to bind and cross epithelial barriers, and that antibodies raised against rH_C_/A W1266L;Y1267S protected against full length BoNT/A at the neuromuscular junction [16], suggesting that the protein was folded and stable. Consistent with this, we observed that the mutant rH_C_/A W1266L;Y1267S protein expresses well and is soluble, similar to the wt rH_C_/A protein (data not shown). In the absence of exogenous GT1b, rH_C_/A showed reduced potency (EC_50_ = 44 nM [95% CI; 36, 54] vs EC_50_ = 27 nM [95% CI; 18, 41]) (overlapping 95% CI) and was less effective at dimerizing FGFR3c than when GT1b had been supplemented, failing to induce complete FGFR3c dimerization even at 200 nM rH_C_/A (**Figure 4; Table 1**). Cells treated with rH_C_/A W1266L;Y1267S in the presence of exogenous GT1b, showed further decreased potency (EC_50_ = 143 nM [95% CI; 127, 161]) (**Figure 4; Table 1**). Interestingly, dimerization of FGFR3c by rH_C_/A in the absence of exogenous GT1b was higher than dimerization by rH_C_/A W1266L;Y1267S, suggesting that rH_C_/A dimerization of FGFR3c utilizes endogenic expressed gangliosides in PC-12 cells. These results suggest that the presence of GT1b and likely additional gangliosides on the cell surface augments rH_C_/A binding and dimerization of the FGFR3 receptor.

**Fig 4.**
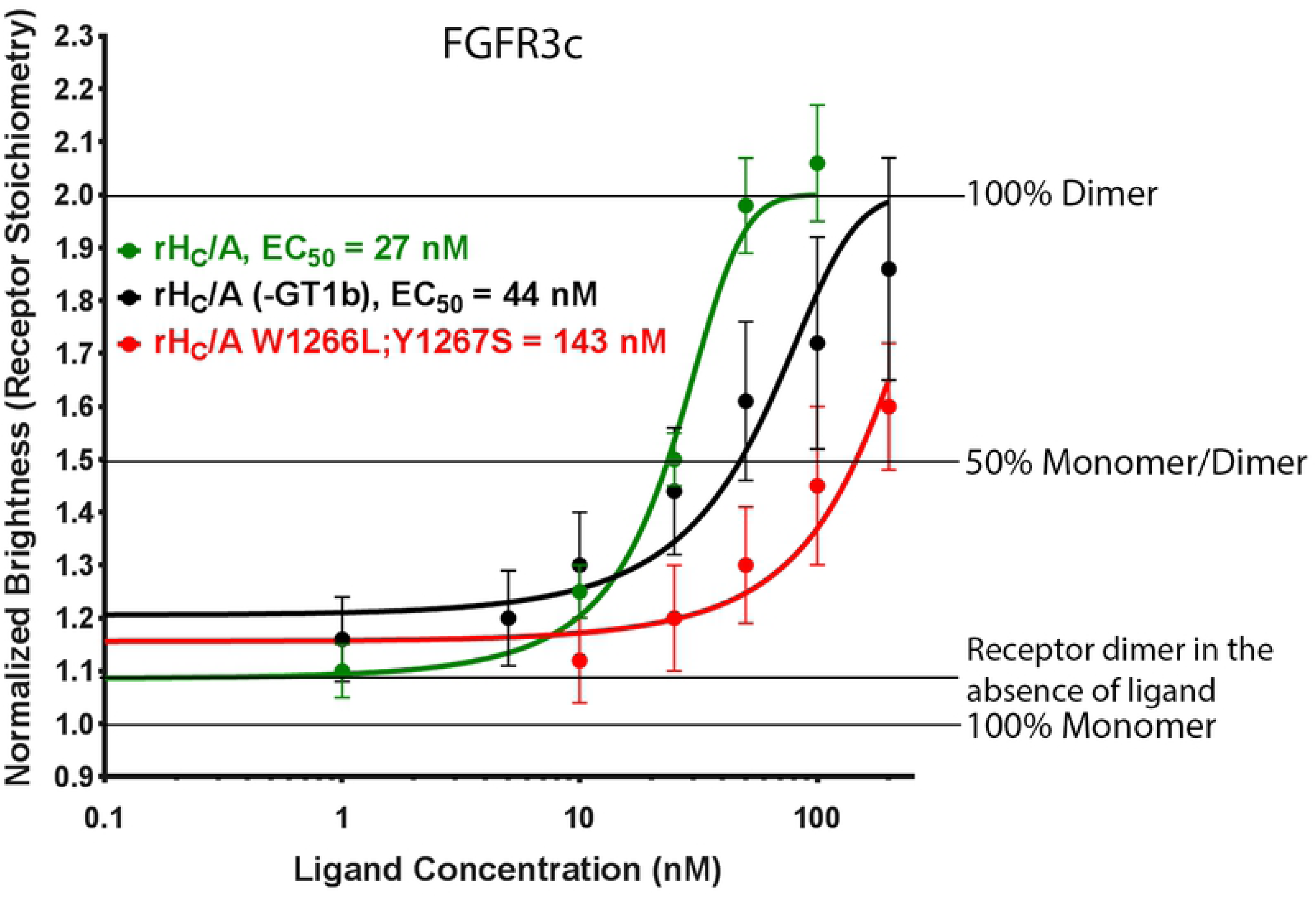
GT1b augments rH_C_/A induced FGFR3c dimerization. BoNT/A binding domain (rH_C_/A) is more potent at dimerizing FGFR3c in the presence of ganglioside (GT1b) (green vs black, EC_50_ = 27 nM [95% CI; 18, 41] vs EC_50_ = 44 nM [95% CI; 36, 54]), and a BoNT/A binding domain (rH_C_/A) ganglioside mutant variant (W1266L;Y1267S) has further reduced ability to dimerize FGFR3c (red) (EC_50_ = 143 nM [95% CI; 127, 161]). Points represent the average normalized brightness values ±SD from greater than 30 cells collected on 4 independent days.

### rH_C_/A dimerizes FGFRs in the rank order FGFR3c > FGFR2b > FGFR1c

As the FGFR subtypes (FGFR1-4) share a high degree of structural and functional homology, it was of interest to assess the affinity of rH_C_/A to other FGFRs, including FGFR1c and FGFR2b (**Figure 5; Table 1**). As expected, ligands known to bind specifically to these receptors (i.e. FGF4/FGFR1c and FGF10/FGFR2b) caused an increase in molecular brightness with complete dimerization appearing around 50-100 nM. The EC_50_ for FGF4 induced FGFR1c dimerization and FGF10 induced FGFR2b dimerization were 17 nM (95% CI; 8, 35) and 19 nM (95% CI; 13, 26), respectively. No increase in molecular brightness was observed with the non-binding receptor ligand pairings FGF10/FGFR1c and FGF4/FGFR2b. Addition of rH_C_/A resulted in a dose-dependent increase in both FGFR1c and FGFR2b dimerization, but with lower potency compared to the native ligands and compared to rH_C_/A-induced FGFR3c dimerization. The EC_50_ for rH_C_/A induced FGFR1c and FGFR2b dimerization were 163 nM (95% CI; 123, 203) and 70 nM (95% CI; 59, 82), respectively.

**Fig 5.**
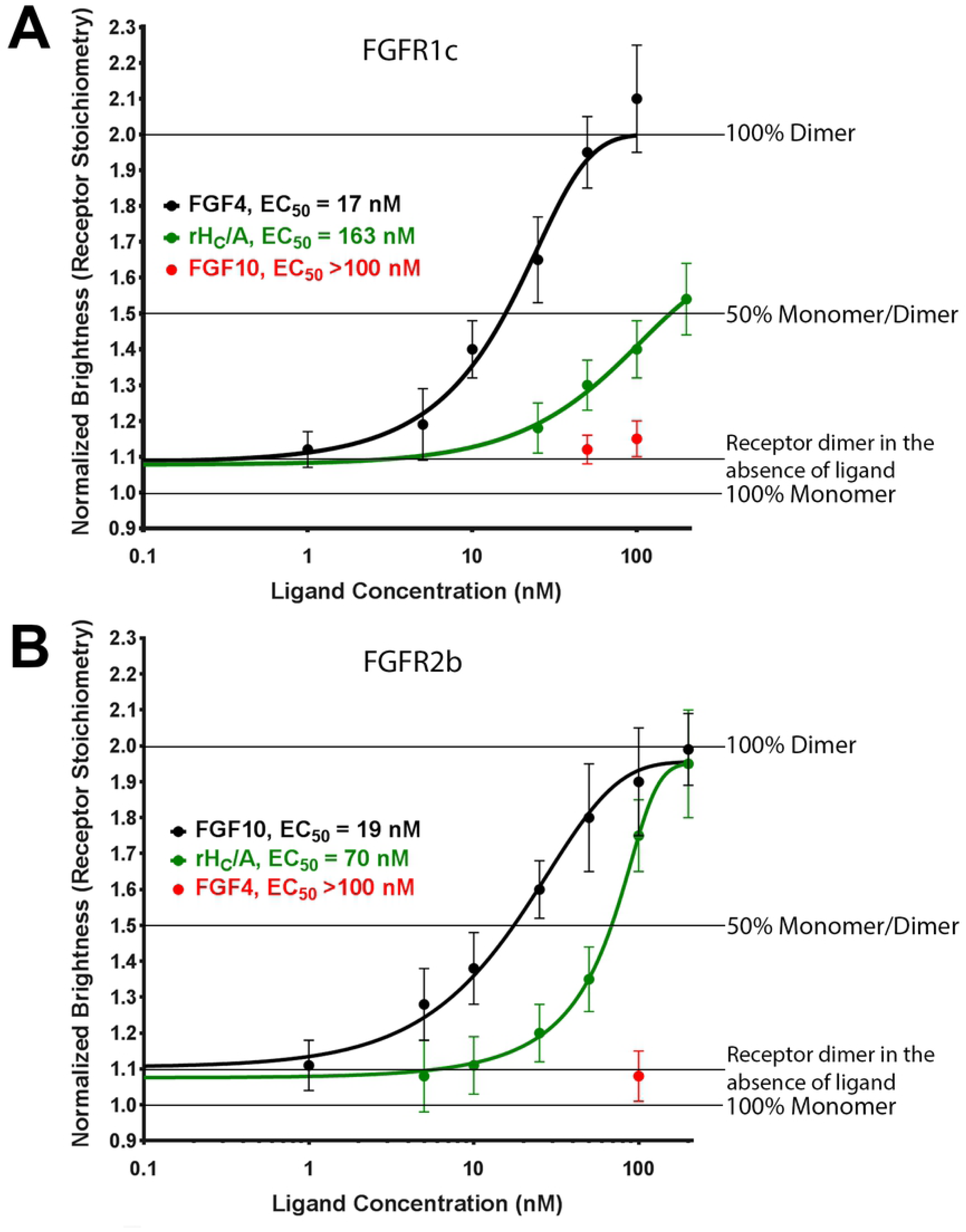
FGFR binding preference of rH_C_/A. **A.** BoNT/A binding domain (rH_C_/A) (green) dimerizes FGFR1c with reduced potency compared to FGFR3c [58]. A known native FGFR1c ligand, FGF4 (black) (positive control), dimerizes FGFR1c (EC_50_ = 17 nM [95% CI; 8, 35]), while a known native FGFR2b ligand, FGF10 (red), does not (EC_50_ >100 nM). Points represent the average normalized brightness values ±SD from greater than 20 cells collected on 4 independent days. **B.** BoNT/A binding domain (rH_C_/A) (green) dimerizes FGFR2b with reduced potency compared to FGFR3c (EC_50_ 70 nM [95% CI; 59, 82] vs 27 nM (95% CI; 18, 41]) (**Figure 3**). A known native FGFR2b ligand, FGF10 (black) (positive control), dimerizes FGFR2b (EC_50_ = 19 nM [95% CI; 13, 26]), while a known native FGFR1c ligand, FGF4 (red), does not (EC_50_ >100 nM). Points represent the average normalized brightness values ±SD from greater than 30 cells collected on 4 independent days. *Based on 2PL with bottom = 1, top = 2.

### rH_C_/A induced dimerization of FGFR3c is dependent on SV2

BoNT/A’s interaction with SV2 has been well characterized and mapped to the fourth luminal domain of SV2 and a beta-strand within H_C_/A [19, 20]. As previous results suggested that FGFR and SV2 interact in cells [24], it was of interest to assess a potential role for SV2 binding during H_C_/A induced FGFR dimerization. Addition of an SV2 mutant variant of H_C_/A (rH_C_/A T1145A;T1146A [20]) resulted in a dose-dependent increase in dimerization of FGFR3c, but with reduced potency (EC_50_ = 78 nM [95% CI; 53, 115]) compared to the wild type variant (EC_50_ = 27 nM [95% CI; 18, 41]) (**Figure 6; Table 1**). Similar to the effect observed with GT1b, these data suggest that residues in H_C_/A which interact with SV2 also affect binding to FGFRs, either directly or indirectly. It is unclear how the T1145A;T1146A mutation affects the folding of rH_C_/A. However, we have observed that the mutant rH_C_/A T1145A;T1146A protein expresses well and is soluble, similar to the wt rH_C_/A protein (data not shown). Previously, it was shown that rH_C_/A T1145A;T1146A bound with lower affinity to a recombinant SV2C luminal domain in vitro [20]. The fact that it still binds, albeit with reduced affinity, also suggests that the protein is folded similarly to the wt protein.

**Fig 6.**
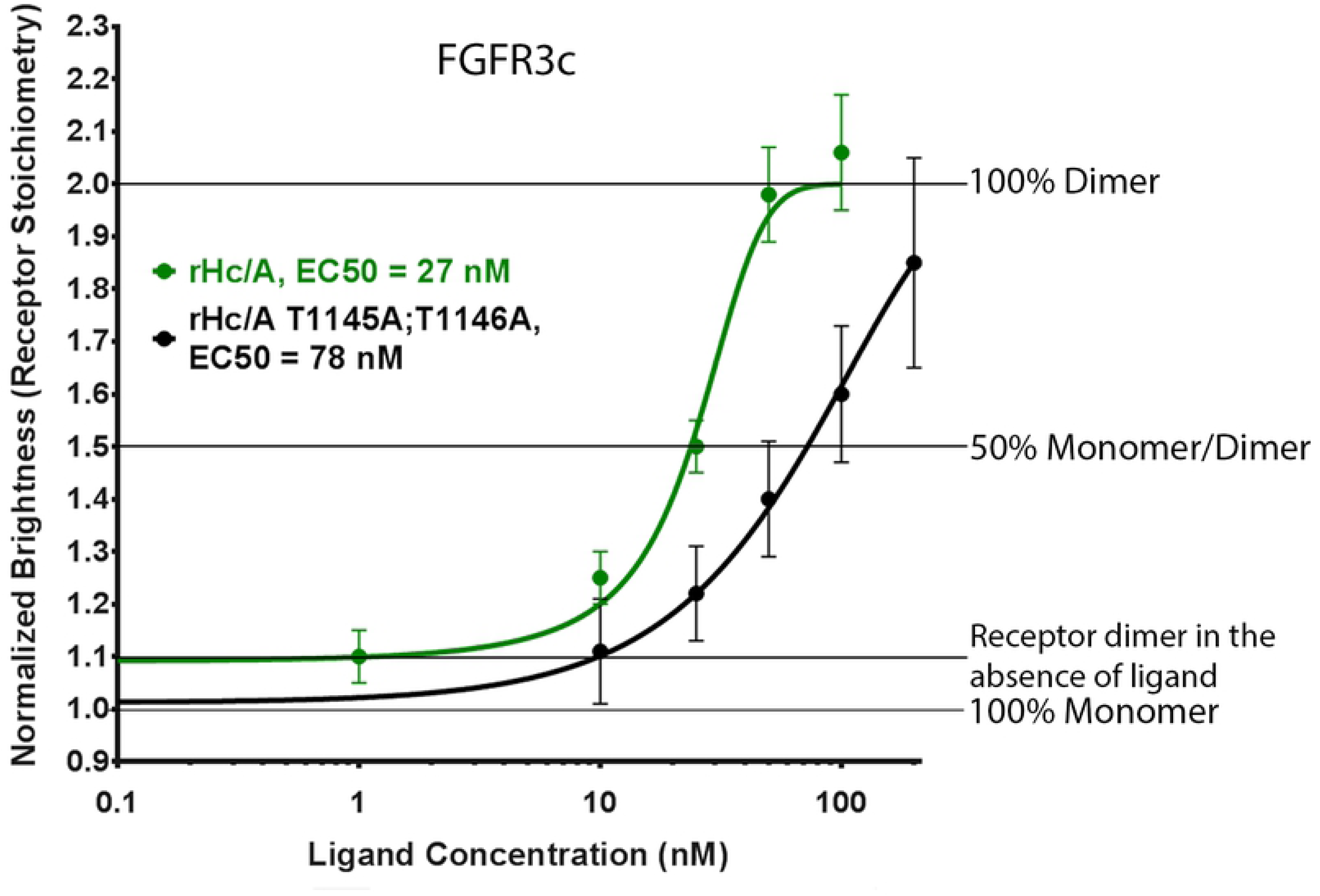
SV2 augments rH_C_/A induced FGFR3c dimerization. A BoNT/A binding domain SV2 mutant variant (rH_C_/A T1145A;T1146A) (black) has reduced ability to dimerize FGFR3c compared to a wt BoNT/A binding domain (rH_C_/A) (green), EC_50_ = 78 nM (95% CI; 53, 115) vs 27 nM (95% CI; 18, 41). Points represent the average normalized brightness values ±SD from greater than 30 cells collected on 4 independent days.

### rH_C_/A does not dimerize EGFR

To assess the selectivity of rH_C_/A for FGFRs versus similar growth factor receptors, the ability of rH_C_/A to dimerize EGFR was assessed. EGFR is another tyrosine kinase growth factor receptor that shares structural and functional homology with FGFRs. The native ligand is EGF [61-63]. As expected, addition of EGF resulted in EGFR dimerization (EC_50_ = 14 nM [95% CI; 11, 19]), while no increase in molecular brightness was observed in the presence of rHc/A, even with upwards of 150 nM (EC_50_ >150 nM) (**Figure S2; Table 1**), suggesting that rH_C_/A has a high degree of specificity for FGFRs.

## DISCUSSION

In the current study, wt and mutant variants of the binding domain of BoNT/A (rH_C_/A) were evaluated in parallel with native FGF ligands possessing known FGFR subtype selectivity to assess H_C_/A’s ability to induce FGFR receptor dimerization. This assessment was done using the FCS & TIRF receptor dimerization assay, a method for measuring molecular associations between proteins by quantifying the molecular brightness of a fluorophore determined through fluctuations in the intensity (due to protein diffusion) within each pixel [49, 53]. This method allows for receptor dynamics to be examined in live cells in response to various ligands, providing continual observation under alternating conditions. The focus on the current investigation was to study FGF receptor dimerization. However, for future studies, this method can potentially provide detailed information on ligand-receptor stoichiometry, cluster formation, and kinetics of receptor endocytosis through the use of fluorescently labeled ligands. The advantage of the FCS & TIRF receptor dimerization assay is that it allows real-time analysis of receptor interactions in live cells as well as direct assessment of the spatial organization of receptor self-association state(s) on the plasma membrane. This presents a new method for addressing biological questions not accessible using conventional terminal endpoint approaches.

Utilizing the FCS & TIRF receptor dimerization assay, dimerization of FGFRs and EGFR were monitored in live neuronal cells in the presence of native and novel receptor ligands, such as the binding domain of BoNT/A, rH_C_/A. Consistent with other reports [64], low level of HaloTag FGF receptor dimerization was observed in the absence of receptor ligand. The analysis was done with cells expressing approximately 100,000-150,000 tagged FGF receptors per cell to minimize potential effects associated with different densities of receptor on the plasma membrane. The observed increase in molecular brightness with increased concentration of ligand was due to dimerization rather than polymerization. This is because the assay exclusively measures the population of receptors with normalized brightness values indicative of monomer or dimer states (**Figure 2**). The fact that a plateau of normalized brightness at ∼2 (or dimer species) was observed for the non-clustering molecular brightness may indicate that these cells have been saturated with ligand to the point that the endocytosis machinery cannot recycle the receptors fast enough to clear dimeric complexes within the assay timeframe.

The EC_50_ values for FGF ligands dimerizing their cognate high affinity receptors ranged from 17-19 nM (95% CI; 8, 35), which is consistent with what has been reported previously for FGFR ligand binding and signaling [24, 35, 39, 65-68], while the EC_50_ values for rH_C_/A were in the rank order FGFR3c (27 nM [95% CI; 18, 41]) > FGFR2b (70 nM [95% CI; 59, 82]) > FGFR1c (163 nM [95% CI; 123, 203]), suggesting that BoNT/A has preferential binding to the FGFR3c receptor subtype. The cellular expression for each receptor subtype was similar (10,000-15,000 counts per cell) and the EC_50_ value for the native ligands was similar for different FGFR subtypes, while the EC_50_ value for H_C_/A differed. Keeping in mind that this is an artificial system, these results support a model where FGFR is part of the specific binding and, potentially, internalization of BoNT/A in neuronal cells. Whether binding of BoNT/A to FGFRs results in pharmacological effects in vivo and in patients is unclear, and it should be kept in mind that clinical toxin doses are in the pM range. Interestingly, unlike most FGFs which are specific to either the b or c isoform of the different receptor subtypes, rH_C_/A dimerized both b and c isoforms, suggesting that H_C_/A has the potential to effect a large number of cell types that express FGFRs [38, 39, 41, 69, 70]. rH_C_/A did not dimerize EGFR at the highest concentration tested here (150 nM) (EC_50_ >150 nM), which indicates rH_C_/A‘s specificity for FGFR over other tyrosine kinase growth factor receptors (summary of the results shown in **Table 1**). Whether FGFRs can facilitate uptake of BoNT/A in other cell types and whether other BoNT serotypes can also bind to FGFRs remains to be further explored.

The observation that gangliosides increase rH_C_/A-mediated dimerization of FGFR3c, while an rH_C_/A variant wherein the residues important for GT1b binding have been mutated (W1266L;Y1267S [16]) showed reduced ability to dimerize, suggests that GT1b and possibly other gangliosides could act as co-receptors for FGFR to facilitate binding to BoNT/A. Although it is unclear whether binding to FGFRs directly triggers internalization of BoNT/A, it is quite likely since binding of FGF native ligands is known to trigger FGFR internalization via mechanisms involving interactions with extended synaptotagmin-like protein, E-Syt2 [71, 72]. This presents a potential evolutionary link between binding of BoNTs to FGFRs and synaptotagmin, the receptor for BoNT/B [73, 74], BoNT/D [75, 76], and BoNT/G [77, 78]. Perhaps BoNT/A binds indirectly to synaptotagmin via its interaction with FGFRs and that the binding of BoNT/A to FGFRs has evolved from an interaction of BoNTs with synaptotagmin.

The specific binding site on H_C_/A for SV2 has been identified as an exposed beta-strand loop in the center of the binding domain [19, 20, 22, 79]. The observation that a variant of H_C_/A (T1145A;T1146A [20]) with mutations in residues important for SV2 binding shows reduced ability to dimerize FGFR3c suggests that these residues, directly or indirectly, affect binding to FGFRs. Perhaps the binding site for FGFR and SV2 overlap and binding of BoNT/A and SV2 is sequential. However, it is also possible that the reduced binding affinity observed by the H_C_/A T1145A;T1146A variant may arise due to perturbations adjacent to the FGFR3c binding region of H_C_/A.

In summary, the data presented here shows that the binding domain of BoNT/A (rH_C_/A) binds to and dimerizes FGFRs in transfected neuronal PC-12 cells. These results further support a model where FGFRs, in particular FGFR3c, which is the primary FGFR subtype expressed in the nervous system [32, 58, 59], function as receptors for BoNT/A and perhaps other BoNT subtypes and serotypes on neuronal cells. The potential pharmacological consequence of BoNT binding to FGFRs, both with respect to BoNT neuronal selectivity and potency and potential receptor mediated effects on neuronal and non-neuronal cell types, is being explored.

## ACKNOWLEDGMENTS

asdfasdfsadfsafds

---

**S1 Figure.**
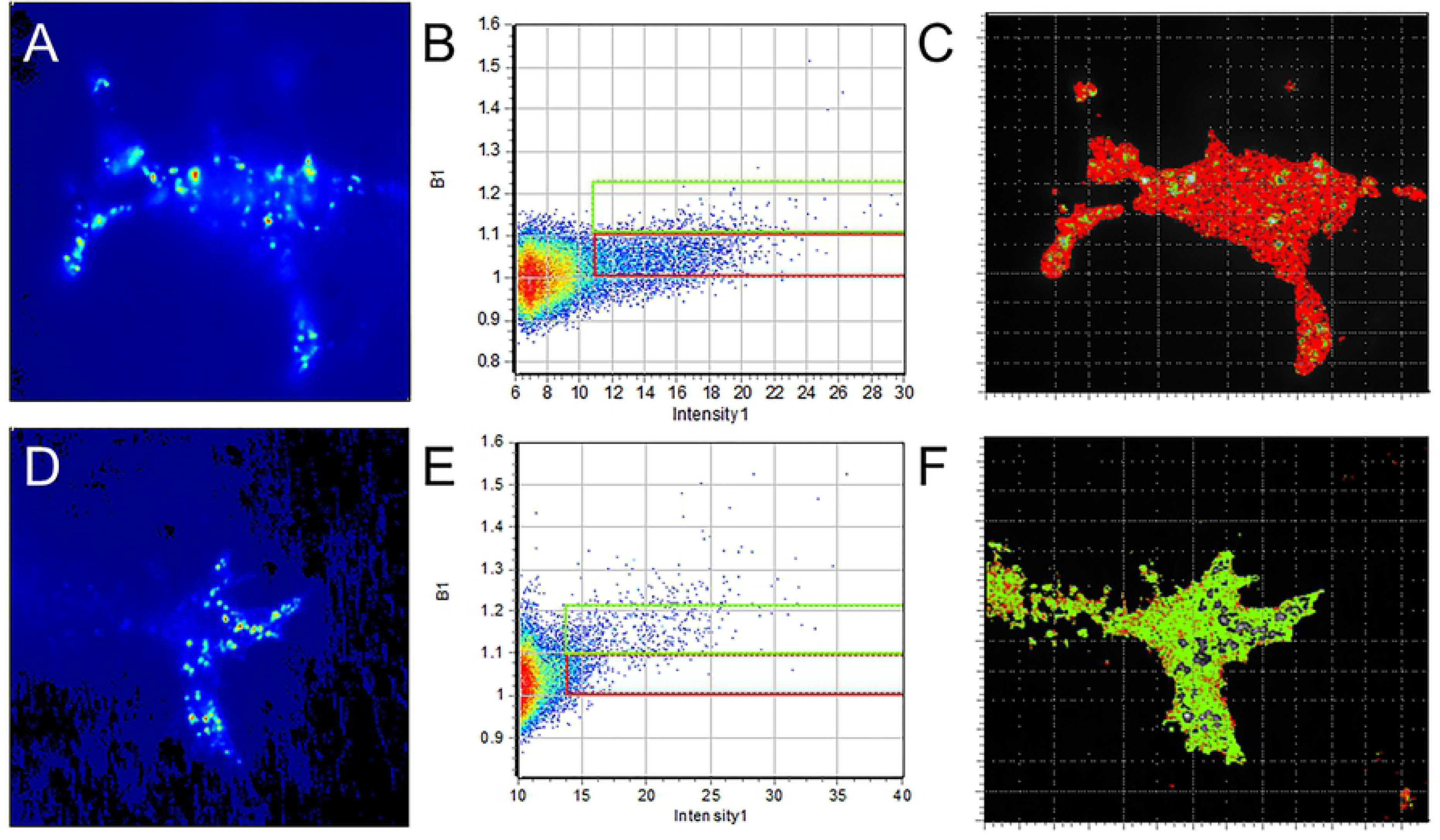
Schematic representation of TIRF N&B analysis of halo tagged FGFR in PC-12 cells. (**A, D**) Intensity images of ∼ 1000 images per cell were collected on cells under (**A**) serum starved conditions followed by treatment with (**D**) growth factor. (**B, E**) The intensity fluctuations within each pixel of the collected images are used to calculate the average intensity and brightness in a 2-D histogram. Regions of interest are selected to determine the population of the monomers (red) and dimers (green). These different populations are represented as an overlay in the images (**C, F**).

**S2 Figure.**
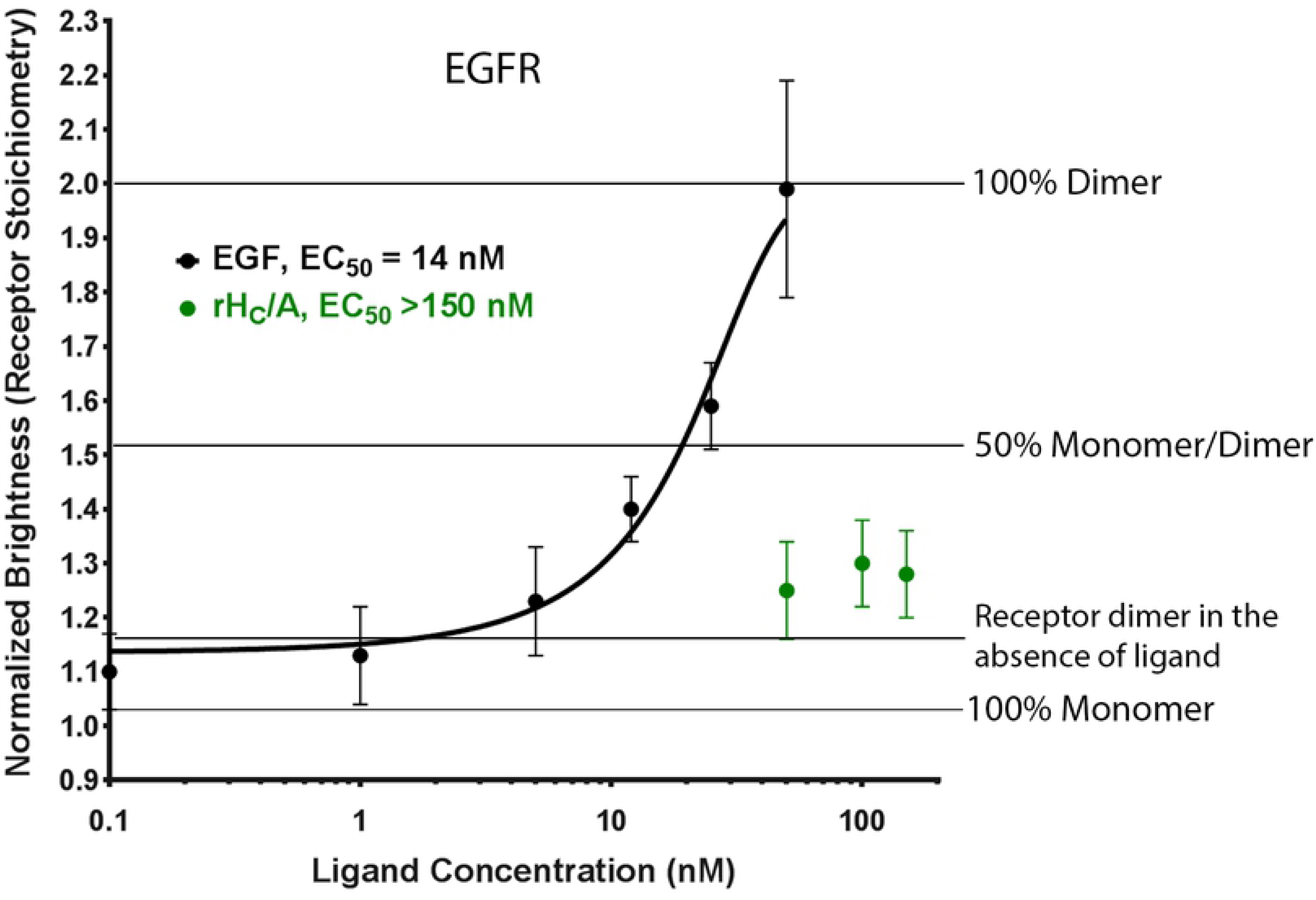
rH_C_/A does not induce EGFR dimerization. BoNT/A binding domain (rH_C_/A) (green) does not dimerize EGFR (EC_50_ >150 nM), while the native ligand for EGFR, EGF (black) can dimerize with high potency (EC_50_ = 14 nM (95% CI; 11, 19]). Points represent the average normalized brightness values ±SD from greater than 30 cells collected on 4 different days.

## REFERENCES

1. Popoff MR, Bouvet P. Clostridial toxins. Future Microbiol. 2009;4(8):1021–64. doi: 10.2217/fmb.09.72. PubMed PMID: 19824793.

2. Rossetto O, Pirazzini M, Montecucco C. Botulinum neurotoxins: genetic, structural and mechanistic insights. Nat Rev Microbiol. 2014;12(8):535–49. doi: 10.1038/nrmicro3295. PubMed PMID: 24975322.

3. Pirazzini M, Rossetto O, Eleopra R, Montecucco C. Botulinum Neurotoxins: Biology, Pharmacology, and Toxicology. Pharmacol Rev. 2017;69(2):200–35. doi: 10.1124/pr.116.012658. PubMed PMID: 28356439; PubMed Central PMCID: PMCPMC5394922.

4. Connan C, Popoff MR. Uptake of Clostridial Neurotoxins into Cells and Dissemination. Curr Top Microbiol Immunol. 2017;406:39–78. doi: 10.1007/82_2017_50. PubMed PMID: 28879524.

5. Monheit GD, Pickett A. AbobotulinumtoxinA: A 25-Year History. Aesthet Surg J. 2017;37(suppl_1):S4–S11. doi: 10.1093/asj/sjw284. PubMed PMID: 28388718; PubMed Central PMCID: PMCPMC5434488.

6. Turton K, Chaddock JA, Acharya KR. Botulinum and tetanus neurotoxins: structure, function and therapeutic utility. Trends Biochem Sci. 2002;27(11):552–8. PubMed PMID: 12417130.

7. Chen S. Clinical uses of botulinum neurotoxins: current indications, limitations and future developments. Toxins (Basel). 2012;4(10):913–39. doi: 10.3390/toxins4100913. PubMed PMID: 23162705; PubMed Central PMCID: PMCPMC3496996.

8. Lacy DB, Tepp W, Cohen AC, DasGupta BR, Stevens RC. Crystal structure of botulinum neurotoxin type A and implications for toxicity. Nat Struct Biol. 1998;5(10):898–902. doi: 10.1038/2338. PubMed PMID: 9783750.

9. Brunger AT, Jin R, Breidenbach MA. Highly specific interactions between botulinum neurotoxins and synaptic vesicle proteins. Cell Mol Life Sci. 2008;65(15):2296–306. doi: 10.1007/s00018-008-8088-0. PubMed PMID: 18425411.

10. Blasi J, Chapman ER, Link E, Binz T, Yamasaki S, De Camilli P, et al. Botulinum neurotoxin A selectively cleaves the synaptic protein SNAP-25. Nature. 1993;365(6442):160–3. doi: 10.1038/365160a0. PubMed PMID: 8103915.

11. Brin MF. Basic and clinical aspects of BOTOX. Toxicon. 2009;54(5):676–82. doi: 10.1016/j.toxicon.2009.03.021. PubMed PMID: 19341758.

12. Aoki KR, Francis J. Updates on the antinociceptive mechanism hypothesis of botulinum toxin A. Parkinsonism Relat Disord. 2011;17 Suppl 1:S28–33. doi: 10.1016/j.parkreldis.2011.06.013. PubMed PMID: 21999893.

13. Yowler BC, Schengrund CL. Botulinum neurotoxin A changes conformation upon binding to ganglioside GT1b. Biochemistry. 2004;43(30):9725–31. doi: 10.1021/bi0494673. PubMed PMID: 15274627.

14. Stenmark P, Dupuy J, Imamura A, Kiso M, Stevens RC. Crystal structure of botulinum neurotoxin type A in complex with the cell surface co-receptor GT1b-insight into the toxin-neuron interaction. PLoS Pathog. 2008;4(8):e1000129. doi: 10.1371/journal.ppat.1000129. PubMed PMID: 18704164; PubMed Central PMCID: PMCPMC2493045.

15. Hamark C, Berntsson RP, Masuyer G, Henriksson LM, Gustafsson R, Stenmark P, et al. Glycans Confer Specificity to the Recognition of Ganglioside Receptors by Botulinum Neurotoxin A. J Am Chem Soc. 2017;139(1):218–30. doi: 10.1021/jacs.6b09534. PubMed PMID: 27958736.

16. Elias M, Al-Saleem F, Ancharski DM, Singh A, Nasser Z, Olson RM, et al. Evidence that botulinum toxin receptors on epithelial cells and neuronal cells are not identical: implications for development of a non-neurotropic vaccine. J Pharmacol Exp Ther. 2011;336(3):605–12. doi: 10.1124/jpet.110.175018. PubMed PMID: 21106906; PubMed Central PMCID: PMCPMC3061530.

17. Rummel A, Mahrhold S, Bigalke H, Binz T. The HCC-domain of botulinum neurotoxins A and B exhibits a singular ganglioside binding site displaying serotype specific carbohydrate interaction. Mol Microbiol. 2004;51(3):631–43. PubMed PMID: 14731268.

18. Mahrhold S, Rummel A, Bigalke H, Davletov B, Binz T. The synaptic vesicle protein 2C mediates the uptake of botulinum neurotoxin A into phrenic nerves. FEBS Lett. 2006;580(8):2011–4. doi: 10.1016/j.febslet.2006.02.074. PubMed PMID: 16545378.

19. Strotmeier J, Mahrhold S, Krez N, Janzen C, Lou J, Marks JD, et al. Identification of the synaptic vesicle glycoprotein 2 receptor binding site in botulinum neurotoxin A. FEBS Lett. 2014;588(7):1087–93. doi: 10.1016/j.febslet.2014.02.034. PubMed PMID: 24583011; PubMed Central PMCID: PMCPMC4067265.

20. Benoit RM, Frey D, Hilbert M, Kevenaar JT, Wieser MM, Stirnimann CU, et al. Structural basis for recognition of synaptic vesicle protein 2C by botulinum neurotoxin A. Nature. 2014;505(7481):108–11. doi: 10.1038/nature12732. PubMed PMID: 24240280.

21. Dong M, Yeh F, Tepp WH, Dean C, Johnson EA, Janz R, et al. SV2 is the protein receptor for botulinum neurotoxin A. Science. 2006;312(5773):592–6. doi: 10.1126/science.1123654. PubMed PMID: 16543415.

22. Yao G, Zhang S, Mahrhold S, Lam KH, Stern D, Bagramyan K, et al. N-linked glycosylation of SV2 is required for binding and uptake of botulinum neurotoxin A. Nat Struct Mol Biol. 2016;23(7):656–62. doi: 10.1038/nsmb.3245. PubMed PMID: 27294781; PubMed Central PMCID: PMCPMC5033645.

23. Weisemann J, Stern D, Mahrhold S, Dorner BG, Rummel A. Botulinum Neurotoxin Serotype A Recognizes Its Protein Receptor SV2 by a Different Mechanism than Botulinum Neurotoxin B Synaptotagmin. Toxins (Basel). 2016;8(5). doi: 10.3390/toxins8050154. PubMed PMID: 27196927; PubMed Central PMCID: PMCPMC4885069.

24. Jacky BP, Garay PE, Dupuy J, Nelson JB, Cai B, Molina Y, et al. Identification of fibroblast growth factor receptor 3 (FGFR3) as a protein receptor for botulinum neurotoxin serotype A (BoNT/A). PLoS Pathog. 2013;9(5):e1003369. doi: 10.1371/journal.ppat.1003369. PubMed PMID: 23696738; PubMed Central PMCID: PMCPMC3656097.

25. Bomba-Warczak E, Vevea JD, Brittain JM, Figueroa-Bernier A, Tepp WH, Johnson EA, et al. Interneuronal Transfer and Distal Action of Tetanus Toxin and Botulinum Neurotoxins A and D in Central Neurons. Cell Rep. 2016;16(7):1974–87. doi: 10.1016/j.celrep.2016.06.104. PubMed PMID: 27498860; PubMed Central PMCID: PMCPMC4988880.

26. Spear PG. Herpes simplex virus: receptors and ligands for cell entry. Cell Microbiol. 2004;6(5):401–10. doi: 10.1111/j.1462-5822.2004.00389.x. PubMed PMID: 15056211.

27. Neal JW, Gasque P. Corrigendum to “The role of primary infection of Schwann cells in the aetiology of infective inflammatory neuropathies” [J Infect 73 (2016) 402-418]. J Infect. 2017;74(5):525. doi: 10.1016/j.jinf.2017.02.003. PubMed PMID: 28245429.

28. Neal JW, Gasque P. The role of primary infection of Schwann cells in the aetiology of infective inflammatory neuropathies. J Infect. 2016;73(5):402–18. doi: 10.1016/j.jinf.2016.08.006. PubMed PMID: 27546064.

29. Tsui CK, Gupta A, Bassik MC. Finding host targets for HIV therapy. Nat Genet. 2017;49(2):175–6. doi: 10.1038/ng.3777. PubMed PMID: 28138150.

30. Dianzani U, Bragardo M, Buonfiglio D, Redoglia V, Funaro A, Portoles P, et al. Modulation of CD4 lateral interaction with lymphocyte surface molecules induced by HIV-1 gp120. Eur J Immunol. 1995;25(5):1306–11. doi: 10.1002/eji.1830250526. PubMed PMID: 7539755.

31. Liu X, Lai C, Wang K, Xing L, Yang P, Duan Q, et al. A Functional Role of Fibroblast Growth Factor Receptor 1 (FGFR1) in the Suppression of Influenza A Virus Replication. PLoS One. 2015;10(4):e0124651. doi: 10.1371/journal.pone.0124651. PubMed PMID: 25909503; PubMed Central PMCID: PMCPMC4409105.

32. Gong SG. Isoforms of receptors of fibroblast growth factors. J Cell Physiol. 2014;229(12):1887–95. doi: 10.1002/jcp.24649. PubMed PMID: 24733629.

33. Yeh BK, Igarashi M, Eliseenkova AV, Plotnikov AN, Sher I, Ron D, et al. Structural basis by which alternative splicing confers specificity in fibroblast growth factor receptors. Proc Natl Acad Sci U S A. 2003;100(5):2266–71. doi: 10.1073/pnas.0436500100. PubMed PMID: 12591959; PubMed Central PMCID: PMCPMC151329.

34. Eswarakumar VP, Lax I, Schlessinger J. Cellular signaling by fibroblast growth factor receptors. Cytokine Growth Factor Rev. 2005;16(2):139–49. doi: 10.1016/j.cytogfr.2005.01.001. PubMed PMID: 15863030.

35. Ornitz DM, Itoh N. The Fibroblast Growth Factor signaling pathway. Wiley Interdiscip Rev Dev Biol. 2015;4(3):215–66. doi: 10.1002/wdev.176. PubMed PMID: 25772309; PubMed Central PMCID: PMCPMC4393358.

36. Beenken A, Mohammadi M. The FGF family: biology, pathophysiology and therapy. Nat Rev Drug Discov. 2009;8(3):235–53. doi: 10.1038/nrd2792. PubMed PMID: 19247306; PubMed Central PMCID: PMCPMC3684054.

37. Belov AA, Mohammadi M. Molecular mechanisms of fibroblast growth factor signaling in physiology and pathology. Cold Spring Harb Perspect Biol. 2013;5(6). doi: 10.1101/cshperspect.a015958. PubMed PMID: 23732477; PubMed Central PMCID: PMCPMC3660835.

38. Chapman JR, Katsara O, Ruoff R, Morgenstern D, Nayak S, Basilico C, et al. Phosphoproteomics of Fibroblast Growth Factor 1 (FGF1) Signaling in Chondrocytes: Identifying the Signature of Inhibitory Response. Mol Cell Proteomics. 2017;16(6):1126–37. doi: 10.1074/mcp.M116.064980. PubMed PMID: 28298517; PubMed Central PMCID: PMCPMC5461542.

39. Zhang X, Ibrahimi OA, Olsen SK, Umemori H, Mohammadi M, Ornitz DM. Receptor specificity of the fibroblast growth factor family. The complete mammalian FGF family. J Biol Chem. 2006;281(23):15694–700. doi: 10.1074/jbc.M601252200. PubMed PMID: 16597617; PubMed Central PMCID: PMCPMC2080618.

40. Liu Y, Ma J, Beenken A, Srinivasan L, Eliseenkova AV, Mohammadi M. Regulation of Receptor Binding Specificity of FGF9 by an Autoinhibitory Homodimerization. Structure. 2017;25(9):1325–36 e3. doi: 10.1016/j.str.2017.06.016. PubMed PMID: 28757146; PubMed Central PMCID: PMCPMC5587394.

41. Degirolamo C, Sabba C, Moschetta A. Therapeutic potential of the endocrine fibroblast growth factors FGF19, FGF21 and FGF23. Nat Rev Drug Discov. 2016;15(1):51–69. doi: 10.1038/nrd.2015.9. PubMed PMID: 26567701.

42. Polanska UM, Fernig DG, Kinnunen T. Extracellular interactome of the FGF receptor-ligand system: complexities and the relative simplicity of the worm. Dev Dyn. 2009;238(2):277–93. doi: 10.1002/dvdy.21757. PubMed PMID: 18985724.

43. Hebert JM. FGFs: Neurodevelopment’s Jack-of-all-Trades – How Do They Do it? Front Neurosci. 2011;5:133. doi: 10.3389/fnins.2011.00133. PubMed PMID: 22164131; PubMed Central PMCID: PMCPMC3230033.

44. Rusnati M, Tanghetti E, Urbinati C, Tulipano G, Marchesini S, Ziche M, et al. Interaction of fibroblast growth factor-2 (FGF-2) with free gangliosides: biochemical characterization and biological consequences in endothelial cell cultures. Mol Biol Cell. 1999;10(2):313–27. PubMed PMID: 9950679; PubMed Central PMCID: PMCPMC25171.

45. Miljan EA, Bremer EG. Regulation of growth factor receptors by gangliosides. Sci STKE. 2002;2002(160):re15. doi: 10.1126/stke.2002.160.re15. PubMed PMID: 12454318.

46. Brewer JR, Mazot P, Soriano P. Genetic insights into the mechanisms of Fgf signaling. Genes Dev. 2016;30(7):751–71. doi: 10.1101/gad.277137.115. PubMed PMID: 27036966; PubMed Central PMCID: PMCPMC4826393.

47. Qian H, Elson EL. Distribution of molecular aggregation by analysis of fluctuation moments. Proc Natl Acad Sci U S A. 1990;87(14):5479–83. PubMed PMID: 2371284; PubMed Central PMCID: PMCPMC54348.

48. James NG, Digman MA, Ross JA, Barylko B, Wang L, Li J, et al. A mutation associated with centronuclear myopathy enhances the size and stability of dynamin 2 complexes in cells. Biochim Biophys Acta. 2014;1840(1):315–21. doi: 10.1016/j.bbagen.2013.09.001. PubMed PMID: 24016602; PubMed Central PMCID: PMCPMC3859711.

49. Unruh JR, Gratton E. Analysis of molecular concentration and brightness from fluorescence fluctuation data with an electron multiplied CCD camera. Biophys J. 2008;95(11):5385–98. doi: 10.1529/biophysj.108.130310. PubMed PMID: 18805922; PubMed Central PMCID: PMCPMC2586556.

50. Jameson DM, Ross JA, Albanesi JP. Fluorescence fluctuation spectroscopy: ushering in a new age of enlightenment for cellular dynamics. Biophys Rev. 2009;1(3):105–18. doi: 10.1007/s12551-009-0013-8. PubMed PMID: 21547245; PubMed Central PMCID: PMCPMC3086800.

51. Jameson DM, James NG, Albanesi JP. Fluorescence fluctuation spectroscopy approaches to the study of receptors in live cells. Methods Enzymol. 2013;519:87–113. doi: 10.1016/B978-0-12-405539-1.00003-8. PubMed PMID: 23280108.

52. James NG, Jameson DM. Steady-state fluorescence polarization/anisotropy for the study of protein interactions. Methods Mol Biol. 2014;1076:29–42. doi: 10.1007/978-1-62703-649-8_2. PubMed PMID: 24108621.

53. Digman MA, Stakic M, Gratton E. Raster image correlation spectroscopy and number and brightness analysis. Methods Enzymol. 2013;518:121–44. doi: 10.1016/B978-0-12-388422-0.00006-6. PubMed PMID: 23276538.

54. Jameson DM. Introduction to Fluorescence. New York, NY: Taylor and Francis; 2014.

55. Ries J, Yu SR, Burkhardt M, Brand M, Schwille P. Modular scanning FCS quantifies receptorligand interactions in living multicellular organisms. Nat Methods. 2009;6(9):643–5. doi: 10.1038/nmeth.1355. PubMed PMID: 19648917.

56. Chen Y, Johnson J, Macdonald P, Wu B, Mueller JD. Observing protein interactions and their stoichiometry in living cells by brightness analysis of fluorescence fluctuation experiments. Methods Enzymol. 2010;472:345–63. doi: 10.1016/S0076-6879(10)72026-7. PubMed PMID: 20580971.

57. Ming AY, Yoo E, Vorontsov EN, Altamentova SM, Kilkenny DM, Rocheleau JV. Dynamics and Distribution of Klothobeta (KLB) and fibroblast growth factor receptor-1 (FGFR1) in living cells reveal the 1. fibroblast growth factor-21 (FGF21)-induced receptor complex. J Biol Chem. 2012;287(24):19997–20006. doi: 10.1074/jbc.M111.325670. PubMed PMID: 22523080; PubMed Central PMCID: PMCPMC3370183.

58. Wuechner C, Nordqvist AC, Winterpacht A, Zabel B, Schalling M. Developmental expression of splicing variants of fibroblast growth factor receptor 3 (FGFR3) in mouse. Int J Dev Biol. 1996;40(6):1185–8. PubMed PMID: 9032024.

59. Fon Tacer K, Bookout AL, Ding X, Kurosu H, John GB, Wang L, et al. Research resource: Comprehensive expression atlas of the fibroblast growth factor system in adult mouse. Mol Endocrinol. 2010;24(10):2050–64. doi: 10.1210/me.2010-0142. PubMed PMID: 20667984; PubMed Central PMCID: PMCPMC2954642.

60. Nagy P, Claus J, Jovin TM, Arndt-Jovin DJ. Distribution of resting and ligand-bound ErbB1 and ErbB2 receptor tyrosine kinases in living cells using number and brightness analysis. Proc Natl Acad Sci U S A. 2010;107(38):16524–9. doi: 10.1073/pnas.1002642107. PubMed PMID: 20813958; PubMed Central PMCID: PMCPMC2944731.

61. Robinson DR, Wu YM, Lin SF. The protein tyrosine kinase family of the human genome. Oncogene. 2000;19(49):5548–57. doi: 10.1038/sj.onc.1203957. PubMed PMID: 11114734.

62. Bakker J, Spits M, Neefjes J, Berlin I. The EGFR odyssey – from activation to destruction in space and time. J Cell Sci. 2017. doi: 10.1242/jcs.209197. PubMed PMID: 29180516.

63. Sarabipour S. Parallels and Distinctions in FGFR, VEGFR, and EGFR Mechanisms of Transmembrane Signaling. Biochemistry. 2017;56(25):3159–73. doi: 10.1021/acs.biochem.7b00399. PubMed PMID: 28621531.

64. Sarabipour S, Hristova K. Mechanism of FGF receptor dimerization and activation. Nat Commun. 2016;7:10262. doi: 10.1038/ncomms10262. PubMed PMID: 26725515; PubMed Central PMCID: PMCPMC4725768.

65. Chen F, Hristova K. The physical basis of FGFR3 response to fgf1 and fgf2. Biochemistry. 2011;50(40):8576–82. doi: 10.1021/bi200986f. PubMed PMID: 21894939; PubMed Central PMCID: PMCPMC3398506.

66. Beenken A, Eliseenkova AV, Ibrahimi OA, Olsen SK, Mohammadi M. Plasticity in interactions of fibroblast growth factor 1 (FGF1) N terminus with FGF receptors underlies promiscuity of FGF1. J Biol Chem. 2012;287(5):3067–78. doi: 10.1074/jbc.M111.275891. PubMed PMID: 22057274; PubMed Central PMCID: PMCPMC3270963.

67. Fernig DG, Rudland PS, Smith JA. Rat mammary myoepithelial-like cells in culture possess kinetically distinct low-affinity receptors for fibroblast growth factor that modulate growth stimulatory responses. Growth Factors. 1992;7(1):27–39. PubMed PMID: 1323979.

68. Ibrahimi OA, Yeh BK, Eliseenkova AV, Zhang F, Olsen SK, Igarashi M, et al. Analysis of mutations in fibroblast growth factor (FGF) and a pathogenic mutation in FGF receptor (FGFR) provides direct evidence for the symmetric two-end model for FGFR dimerization. Mol Cell Biol. 2005;25(2):671–84. doi: 10.1128/MCB.25.2.671-684.2005. PubMed PMID: 15632068; PubMed Central PMCID: PMCPMC543411.

69. Turner N, Grose R. Fibroblast growth factor signalling: from development to cancer. Nat Rev Cancer. 2010;10(2):116–29. doi: 10.1038/nrc2780. PubMed PMID: 20094046.

70. Nunes QM, Li Y, Sun C, Kinnunen TK, Fernig DG. Fibroblast growth factors as tissue repair and regeneration therapeutics. PeerJ. 2016;4:e1535. doi: 10.7717/peerj.1535. PubMed PMID: 26793421; PubMed Central PMCID: PMCPMC4715458.

71. Jean S, Tremblay MG, Herdman C, Guillou F, Moss T. The endocytic adapter E-Syt2 recruits the p21 GTPase activated kinase PAK1 to mediate actin dynamics and FGF signalling. Biol Open. 2012;1(8):731–8. doi: 10.1242/bio.2012968. PubMed PMID: 23213466; PubMed Central PMCID: PMCPMC3507230.

72. Jean S, Mikryukov A, Tremblay MG, Baril J, Guillou F, Bellenfant S, et al. Extended-synaptotagmin-2 mediates FGF receptor endocytosis and ERK activation in vivo. Dev Cell. 2010;19(3):426–39. doi: 10.1016/j.devcel.2010.08.007. PubMed PMID: 20833364.

73. Dong M, Richards DA, Goodnough MC, Tepp WH, Johnson EA, Chapman ER. Synaptotagmins I and II mediate entry of botulinum neurotoxin B into cells. J Cell Biol. 2003;162(7):1293–303. doi: 10.1083/jcb.200305098. PubMed PMID: 14504267; PubMed Central PMCID: PMCPMC2173968.

74. Atassi MZ, Taruishi M, Naqvi M, Steward LE, Aoki KR. Synaptotagmin II and gangliosides bind independently with botulinum neurotoxin B but each restrains the other. Protein J. 2014;33(3):278–88. doi: 10.1007/s10930-014-9557-y. PubMed PMID: 24740609.

75. Peng L, Berntsson RP, Tepp WH, Pitkin RM, Johnson EA, Stenmark P, et al. Botulinum neurotoxin D-C uses synaptotagmin I and II as receptors, and human synaptotagmin II is not an effective receptor for type B, D-C and G toxins. J Cell Sci. 2012;125(Pt 13):3233–42. doi: 10.1242/jcs.103564. PubMed PMID: 22454523; PubMed Central PMCID: PMCPMC4067266.

76. Berntsson RP, Peng L, Svensson LM, Dong M, Stenmark P. Crystal structures of botulinum neurotoxin DC in complex with its protein receptors synaptotagmin I and II. Structure. 2013;21(9):1602–11. doi: 10.1016/j.str.2013.06.026. PubMed PMID: 23932591; PubMed Central PMCID: PMCPMC3803103.

77. Rummel A, Karnath T, Henke T, Bigalke H, Binz T. Synaptotagmins I and II act as nerve cell receptors for botulinum neurotoxin G. J Biol Chem. 2004;279(29):30865–70. doi: 10.1074/jbc.M403945200. PubMed PMID: 15123599.

78. Schmitt J, Karalewitz A, Benefield DA, Mushrush DJ, Pruitt RN, Spiller BW, et al. Structural analysis of botulinum neurotoxin type G receptor binding. Biochemistry. 2010;49(25):5200–5. doi: 10.1021/bi100412v. PubMed PMID: 20507178; PubMed Central PMCID: PMCPMC2894633.

79. Mahrhold S, Bergstrom T, Stern D, Dorner BG, Astot C, Rummel A. Only the complex N559-glycan in the synaptic vesicle glycoprotein 2C mediates high affinity binding to botulinum neurotoxin serotype A1. Biochem J. 2016;473(17):2645–54. doi: 10.1042/BCJ20160439. PubMed PMID: 27313224.

80. Chen Y, Muller JD, So PT, Gratton E. The photon counting histogram in fluorescence fluctuation spectroscopy. Biophys J. 1999;77(1):553–67. doi: 10.1016/S0006-3495(99)76912-2. PubMed PMID: 10388780; PubMed Central PMCID: PMCPMC1300352.

